# Knowledge and awareness of cervical cancer in Southwestern Ethiopia is lacking: a descriptive analysis

**DOI:** 10.1101/592196

**Authors:** Atif Saleem, Alemayehu Bekele, Megan B. Fitzpatrick, Eiman A. Mahmoud, Athena W. Lin, H. Eduardo Velasco, Mona M. Rashed

## Abstract

**Purpose:** Cervical cancer remains the second most common cancer and cancer-related death among women in Ethiopia. This is the first study, to our knowledge, describing the demographic, and clinicopathologic characteristics of cervical cancer cases in a mainly rural, Southwestern Ethiopian population with a low literacy rate to provide data on the cervical cancer burden and help guide future prevention and intervention efforts.

**Methods:** A descriptive analysis of 154 cervical cancer cases at the Jimma University Teaching Hospital in Southwestern Ethiopia from January 2008 – December 2010 was performed. Demographic and clinical characteristics were obtained from patient questionnaires and cervical punch biopsies were histologically examined.

**Results:** Of the 154 participants with a histopathologic diagnosis of cervical cancer, 95.36% had not heard of cervical cancer and 89.6% were locally advanced at the time of diagnosis. Moreover, 86.4% of participants were illiterate, and 62% lived in a rural area.

**Conclusion:** A majority of the 154 women with cervical cancer studied at the Jimma University Teaching Hospital in Southwestern Ethiopia were illiterate, had not heard of cervical cancer and had advanced disease at the time of diagnosis. Given the low rates of literacy and knowledge regarding cervical cancer in this population which has been shown to correlate with a decreased odds of undergoing screening, future interventions to address the cervical cancer burden here must include an effective educational component.

## Introduction

Cervical cancer pathology and demographic data is lacking from Southwestern Ethiopia. The Jimma University Teaching Hospital (JUTH) is located in the city of Jimma which is 352 km southwest of Ethiopia’s capital city Addis Ababa and is unique in that it acts as the only teaching and referral hospital in the region, serving a population of 15 million people [1]. Moreover, Jimma is part of the Oromia state which has one of the highest poverty rates (74.9% of the population) and lowest literacy rates in the country (36% of all residents, with 17% of the female residents living in rural settings) [2-3]. Contributory data from this hospital is vital since every year, an estimated 7,095 women are diagnosed with cervical cancer and 4,732 deaths are due to the disease in Ethiopia - it is currently the second most common cause of female cancer deaths in Ethiopia, after breast cancer.

Infection with high-risk human papillomavirus (HPV) is the necessary cause of >99% of cervical cancer [4]. Other contributing factors include smoking, total fertility rate, and human immunodeficiency virus (HIV) infection [5]. The knowledge about cervical cancer in Ethiopia has been reported to range from 21.2% to 53.7%, with screening rates that ranged from 9.9% to 23.5%. Three of these four studies, however, took place in Northern Ethiopia [6-9]. Though there is not yet an organized cervical cancer education or screening program in Ethiopia, the ongoing dilemma remains how much the absence of such programs compared to a general lack of education or negative attitude towards cervical cancer contribute to the disease burden. Aweke et al. described that 34.8% of 583 survey respondents in Southern Ethiopia had a negative attitude pertaining to cervical cancer [7].

## Materials and methods

### Place of study

The study took place at the Jimma University Teaching Hospital Departments of Obstetrics and Gynecology and Medical Laboratory Sciences and Pathology in Southwestern Ethiopia. This study was approved by the Touro University California Institutional Review Board in the United States of America, by the Research and Publication Committee of the Faculty of Medical Sciences at Jimma University, by the Jimma University Ethics Review Committee and by the Jimma University Teaching Hospital Departments of Obstetrics and Gynecology and Medical Laboratory Sciences and Pathology in Ethiopia.

### Study population

The study population included non-pregnant women voluntarily attending the Jimma University Teaching Hospital Department of Obstetrics and Gynecology outpatient clinic from January 2008 – December 2010 who had evidence of cervical lesions on initial pelvic examination. All of the participants voluntarily presented to the clinic and were willing to be screened; data was collected only after full informed oral consent for participating in the study was obtained.

### Screening procedure

Data was collected by residents in the Department of Obstetrics and Gynecology who were informed regarding the study parameters and were in charge of the outpatient service on a rotation basis. All non-pregnant women with cervical lesions were invited to participate during the study time period. The patients were informed about the indications, contraindications, and alternative options of undergoing a cervical punch biopsy to recognize any cervical pathology. Oral consent was obtained from each case before the interview, punch biopsy procedure and data collection for participation in the study. Then each patient was interviewed using a standardized questionnaire to extract information regarding additional clinical features, sociodemographic characteristics, maternity history, and knowledge about cervical carcinoma, amongst others. Questionnaires were collected weekly and checked for adequacy - those with inadequate data (missing data or unrecognizable responses) were excluded. Pelvic examination was conducted to characterize the cervical lesion(s) and determine the clinical stage. Thorough speculum examination of the cervix was performed to describe any lesion(s) and subsequently a four quadrant punch biopsy of the cervix was taken. The biopsy material was preserved in 10% formaldehyde and submitted to the Department of Medical Laboratory Sciences and Pathology.

In the Department of Pathology the formalin fixed tissue was embedded in paraffin, sections were cut and subsequently stained as described. From each case, four microscopic slides were prepared – one remained in the Department of Pathology for clinical management and three were used for the current study. The slide used for clinical management was stained with hematoxylin and eosin (H&E) and diagnosed by a pathologist in the Department of Pathology according to the World Health Organization histological classification of tumors of the uterine cervix and this pathologic report was recorded and relayed to the physician specific to the case for clinical care. The H&E study slides were identified by the biopsy and code number assigned by the initial physician on the biopsy request sheet and questionnaire and were submitted for diagnosis to a pathologist from Touro University California who was blinded regarding the case for quality control. If there was disagreement in the reports between the slide used for clinical management and the second observer report, the slide was given to a third pathologist and the agreement of the two pathologists was taken as the gold standard report to be recorded.

## Data Analysis

Data was initially entered into Microsoft Excel after which it was coded and analyzed using STATA 15.0 software. Data cleaning was performed only in the form of eliminating missing data so as to improve accuracy, and descriptive statistics were subsequently used to summarize all variables.

## Results

A total of 240 women presented with various gynecological complaints to the outpatient clinic from January 2008 – December 2010. Eighty six women were excluded: 30 of these women had a diagnosis other than cervical cancer such as cervicitis or a cervical polyp but their remaining data was insufficient to analyze; the remaining 56 women were excluded due to an uninterpretable or equivocal biopsy. This left 154 cases to be analyzed and their subjective and objective clinical data is summarized in Tables 1 - 3.

**Table 1.** Selected demographic and clinical features of 154 cervical cancer cases at the Jimma University Teaching Hospital, Ethiopia from January 2008 – December 2010.

| Variable | Mean | Standard Deviation | Number Missing |
| --- | --- | --- | --- |
| Age | 45.19 | 11.17 | 1 (0.6%) |
| Parity | 6.27 (median = 6) | 2.505 (IQR = 3) | 3 (1.94%) |
| Lifetime number of sexual partners | 2.909 (median = 1) | 6.53 (IQR = 1) | 0 (0%) |
| Age (in years) at first intercourse | 15.83 | 2.08 | 16 (10.38%) |
| Age (in years) at first pregnancy | 17.88 | 2.38 | 13 (8.44%) |
| Time (in months) of irregular vaginal bleeding | 5.78 (median = 3) | 8.10 (IQR = 5.42) | 27 (17.53%) |
| Time (in months) of Post-Coital Bleeding | 6.15 (median = 4) | 5.56 (IQR = 7.62) | 96 (62.33%) |
| Time (in months) of Vaginal Discharge | 5.27 (median = 3) | 5.18 (IQR = 5) | 61 (39.61%) |
| Time (in months) of Pelvic Pain | 4.96 (median = 3) | 5.63 (IQR = 5) | 69 (44.805%) |

IQR: Interquartile range

**Table 2.** Selected non-quantifiable demographic and clinical features of 154 cervical cancer cases at the Jimma University Teaching Hospital, Ethiopia from January 2008 – December 2010.

| Category | N (%) | Missing (%) |
| --- | --- | --- |
| Ethnicity |  | 1 (0.6%) |
| Oromo | 105 (68.6%) |  |
| Amhara | 20 (13.07%) |  |
| Other<sup>a</sup> | 28 (18.3%) |  |
| Religion |  | 4 (2.59%) |
| Muslim | 101 (67.33%) |  |
| Orthodox | 41 (27.33%) |  |
| Protestant | 8 (5.33%) |  |
| Marital Status |  | 6 (3.89%) |
| Married | 107 (72.29%) |  |
| Widowed | 22 (14.86%) |  |
| Divorced | 16 (10.81%) |  |
| Single | 3 (2.02%) |  |
| Smoking |  | 1 (0.6%) |
| No | 151 (96.69%) |  |
| Yes | 2 (1.3%) |  |
| Contraception<sup>b</sup> |  | 4 (2.59%) |
| No | 115 (76.66%) |  |
| Yes | 35 (23.33%) |  |
| HIV Status (self-reported) |  | 108 (72%) |
| Negative | 42 (91.3%) |  |
| Positive | 4 (8.69%) |  |
| Heard of Cervical Cancer |  | 3 (1.94%) |
| No | 144 (95.36%) |  |
| Yes<sup>c</sup> | 7 (4.63%) |  |
| Partner Circumcised? |  | 24 (15.58%) |
| Yes | 120 (92.3%) |  |
| No | 10 (7.69%) |  |
| Illiterate |  |  |
| Yes | 133 (86.4%) |  |
| No | 21 (13.6%) |  |
| Lived in Rural Location |  |  |
| Yes | 95 (62%) |  |
| No | 59 (38%) |  |

<sup>a</sup>Other ethnicities included Shekicho, Gurage, Kulo, Yem, Kefa, Dawro, and Bench.
<sup>b</sup>Amongst those that admitted to using contraception, none practiced barrier contraception- only oral contraceptive pills or injectable contraceptives were used.
<sup>c</sup>Out of those that have heard of cervical cancer, all denied knowing the cause of it.

**Table 3.** Selected objective clinical features of 154 cervical cancer cases at the Jimma University Teaching Hospital, Ethiopia from January 2008 – December 2010.

| Category | N (%) | Missing (%) |
| --- | --- | --- |
| Cervical Lesion Appearance<br>on Pelvic Exam |  | 3 (1.94%) |
| Fungating | 90 (59.6%) |  |
| Diffuse/Infiltrating | 33 (21.85%) |  |
| Ulcerating | 22 (14.56%) |  |
| Polyp | 6 (3.97%) |  |
| Stage |  | 4 (2.59%) |
| I | 12 (8%) |  |
| IIA | 32 (21.33%) |  |
| IIB | 47 (31.33%) |  |
| IIIA | 33 (22%) |  |
| IIIB | 18 (12%) |  |
| IVA | 6 (4%) |  |
| IVB | 2 (1.33%) |  |
| Cervical Carcinoma Subtype |  | 0 (0%) |
| Keratinizing Squamous<br>Cell Carcinoma | 79 (51.29%) |  |
| Large Cell Non-Keratinizing Squamous Cell<br>Carcinoma | 59 (38.31%) |  |
| Small Cell Carcinoma | 9 (5.84%) |  |
| Adenocarcinoma | 4 (2.59%) |  |
| Basaloid Squamous Cell<br>Carcinoma | 2 (1.298%) |  |
| Adenosquamous Carcinoma | 1 (0.649%) |  |

## Discussion

### Demographic and clinical features

Cervical cancer is a unique cancer in that effective screening methods are known to prevent disease and associated mortality. Knowledge about the disease and preventive options are vital to effectively control the disease; however, we highlight in the current study that there is a considerable lack of knowledge and awareness regarding cervical cancer which is the second most common cause of cancer deaths in Ethiopia.

Knowledge about cervical cancer in Ethiopia has been reported to range from 21.2% to 53.7% [6-9], and Aweke et. Al described that 34.8% (n=583) of survey respondents in Southern Ethiopia had a negative attitude pertaining to cervical cancer [7]. In our study a majority 144 women (95.36%) had not heard of cervical cancer compared to 138 out of 633 women (21.8%) who had not heard of it in a study done in Gondar town, northwest Ethiopia in 2010 [6]. In that cross-sectional survey, the literacy rate was 18.8%, whereas the rate was 86.4% in our current study. Moreover, a majority of our study participants lived in rural areas (62%) where access to television/radio and health professionals is limited-these were noted as the two most common sources for hearing about cervical cancer in the aforementioned study. The lack of knowledge regarding cervical cancer is of note since preventative efforts such as screening have been shown to reduce the risk of cervical cancer compared to no screening [10]; furthermore, a single-visit approach for cervical cancer screening in Ethiopia was described by Addis Tesfa in 2010 where visual inspection of the cervix with acetic acid wash (VIA) with subsequent cryotherapy of premalignant lesions was performed. One VIA at age 35 can reduce a woman’s lifetime risk of cervical cancer by 25% and if screened again at age 40 by 65% [11].

Cervical cancer educational strategies have been shown to improve screening in studies which targeted rural populations of sub-Saharan Africa [12-14]. Erku et al. describe that the odds of undergoing cervical cancer screening among women who had a comprehensive knowledge on cervical cancer and screening were 2.02 times higher than those who did not in a northwest Ethiopian population [8]. In this study, a majority (87.7%) of the respondents had heard of cervical cancer. This is likely an overestimate since this study included a population of women living with HIV/acquired immunodeficiency syndrome (AIDS) which may have an increased level of awareness with more frequent healthcare visits.

In Ethiopia, currently there are approximately 25 cervical cancer screening centers that are providing visual inspection with acetic acid (VIA), however there is low participation in the community which is partly attributed to the lack of awareness regarding this disease [15]. Geremew et al describe that college and above educational status, knowing someone with cervical cancer, and having knowledge of cervical cancer were positively associated with favorable attitudes towards cervical cancer screening [16]; in the current study, a majority of the patients were illiterate and had decreased knowledge regarding cervical cancer which may explain the lack of screening in our specific population. The National Cancer Control Plan of Ethiopia headed by the Federal Ministry of Health Ethiopia plans a nation-wide scale up of the screening and treatment of cervical pre-cancerous lesions into over 800 health facilities [17]. The mean age at diagnosis of cervical cancer in the United States has been shown to be 48 years and in our study from Ethiopia it was 45 years [18]. Our study differs in that there is no data on prior screening which may have decreased the age at diagnosis and if so, could be attributed to a possible faster progression from HPV to cervical cancer secondary to HIV co-infection or other synergistic risk factors, particularly in the absence of a cervical cancer screening program. Established risk factors for most cervical cancer include: early onset of sexual activity, multiple sexual partners immunosuppression, increasing parity, low socioeconomic status and oral contraceptive use [5].

A qualitative study of 198 patients with cervical cancer from Tikur Anbessa Hospital in Addis Ababa, Ethiopia in 2013 [19] is compared to our study at JUTH in Table 4. The mean age at first sexual intercourse in southwestern Ethiopia has previously been shown to be 17.07 years (+/- 2.12) in a group of 405 young women where cervical lesions were not studied [20]. Our data of cervical cancer cases shows the mean age at first sexual intercourse to be 15.83 years (+/- 2.08) and the mean age from the Tikbur Anbessa study is 16.5 years which may be explained by the cultural practice of marriage at a younger age in these selected populations.

**Table 4.** Comparison of data pertinent to selected risk factors for cervical cancer from Jimma University Teaching Hospital in southwestern Ethiopia (January 2008 – December 2010) and Tikbur Anbessa Hospital in Addis Ababa, Ethiopia (April 2013) [19] from women with a diagnosis of cervical cancer to that of representative women in Ethiopia (where cervical lesions were not necessarily studied).

| Variable | Jimma University Teaching Hospital | Tikur Anbessa Hospital | Ethiopia (cervical lesions not studied) |
| --- | --- | --- | --- |
| Mean age (in years) at first sexual intercourse | 15.83 | 16.5 | 17.07 |
| Mean number of sexual partners | 2.9 | 1.6<sup>a</sup> | 1.4 |
| Parity | 6.27 | 5.3 | 4.8 |

<sup>a</sup>(Out of 198 respondents, 52.3% responded 1, 33% responded 2 and 29% responded 3 or more).

Prior studies found that the mean number of sexual partners in Ethiopia for women is approximately 1.5 (cervical lesions not specified) compared to our study which is 2.9 [21-22] and an increased number of sexual partners raises the probability of becoming infected with HPV. The total fertility rate is estimated to be 4.8 children per woman in Ethiopia (cervical lesions not specified) compared to our study which is 6.27 per woman. The proposed mechanism for higher parity as a risk factor for cervical cancer include increased estrogen exposure during pregnancy, persistence of the transformation zone on the ectocervix in multiparous women, and cervical tissue damage during vaginal deliveries [22].

Hormonal steroids (such as those in oral contraceptive pills) have been shown to activate enhancer elements in the upstream regulatory region of the HPV type 16 viral genome which is one proposed mechanism for the increased risk of cervical cancer [23]. Out of the 35 women (23.33%) in our study used contraception, none practiced barrier contraception. The majority of these 35 women (80%) used oral contraceptive pills which have been shown to increase the cumulative incidence of invasive cervical cancer by age 50 from 7.3 to 8.3 per 1000 in developing countries [24].

This study took place during the rapid expansion phase of HIV/AIDS services in Ethiopia where the number of patients on antiretroviral therapy (ART) increased from 900 at the beginning of 2005 to over 150,000 by June 2008 [25]. Despite this increase in ART use, the frequency of cervical cancer cases in Ethiopia has increased from 2005 until present, with a yearly increment from 1997-2012 except in 1999 and 2009 [26]. This increase may, however, be attributed to increased awareness, screening and subsequent diagnosis. In our study, a majority of women presented at stage IIB followed by stage IIIA at the time of diagnosis and the general trends in Ethiopia at that time remained at presenting at stage IIIB being the most frequent, and secondly stage IIB (Table 5).

**Table 5.** Clinical stages of cervical carcinoma cases.

| Clinical Stage | N (%) |
| --- | --- |
| I | 12 (8%) |
| IIA | 32 (21.33%) |
| IIB | 47 (31.33%) |
| IIIA | 33 (22%) |
| IIIB | 18 (12%) |
| IVA | 6 (4%) |
| IVB | 2 (1.33%) |

## Histopathologic classification

The majority of cervical cancers in the United States are squamous cell carcinoma (69%) followed by adenocarcinoma (25%) [27]. Histopathologic subtype classification in a study of 598 cervical cancer cases in Nigeria and 2,930 cervical cancer cases in South Africa demonstrated squamous cell carcinoma as the most common type as was shown in 92.3% and greater than 80% of cases, respectively [28-29]. In the United States, other non-squamous cervical cancers have been observed in the following frequencies: adenosquamous carcinomas represent 20%-30% of all adenocarcinomas of the cervix and small cell carcinomas represent 0.5%-5% of all invasive cervical cancers. In our study, approximately 91% of the cervical cancer cases were squamous cell carcinomas (including keratinizing, non-keratinizing and basaloid subtypes), 5.84% were small cell carcinomas, 2.59% were adenocarcinomas, and 0.64% were adenosquamous carcinomas. The squamous cell carcinoma frequency was similar to that observed in prior studies; however, an increased frequency of small cell carcinomas over adenocarcinomas was also noted in our study. It has been shown that the keratinizing squamous cell carcinoma subtype is associated with a higher likelihood of advanced stage disease and a lower overall 5-year survival [30] and in our study we observed a 51.29% frequency of this subtype.

The HPV-18 genotype is more commonly associated with adenocarcinomas and small cell carcinomas of the cervix; however, the cases in this study were not subtyped. Few studies describing the high-risk HPV genotypes have been performed in Ethiopia out of which one study of 98 women with cervical dysplasia in Jimma showed that HPV-18 was detected in 8.2% of the 67.1% of HPV DNA positive samples [31]. Based on other studies, HPV type 18 is detected in 18.2% of cervical cancer cases in Ethiopia [32].

A population based study from 1988-2004 of 6,853 women with squamous cell carcinoma found that keratinizing squamous cell carcinoma of the cervix may be less radiosensitive and associated with shorter overall survival than non-keratinizing squamous cell carcinoma [30]. In our study, a majority of women presented with locally advanced cervical cancer (89.6%, Table 1), whereas approximately 54.9-58.8% of patients were diagnosed at a late stage in a California database from the United States [33], as a means of comparison to a high-income country with an established screening program in place. We believe the majority of women in our study presented with locally advanced lesions not entirely due to an intrinsic pathogenetic difference, but because of lack of a cervical screening program in Ethiopia, decreased knowledge about cervical cancer, inability to attend health clinics due to cost and travel expenditure, and increased exposure to risk factors.

### Limitations, future directions and recommendations

Our study did not perform laboratory confirmation of HPV or HIV infection, or test for co-infections with other sexually transmitted infections. Recall bias may have affected the demographic data since it was procured by a survey. Future directions include measuring survival outcomes after intervention for cervical cancer and studying the effectiveness of cervical cancer screening after education. Based on our data, in this specific population of Ethiopian women we recommend promoting an educational initiative about cervical cancer among Ethiopian women given that improved knowledge regarding the disease has been shown to increase screening and decrease cervical cancer rates.

## Conclusions

Most of the 154 women with cervical cancer studied at the JUTH in southwestern Ethiopia were illiterate, had not heard of cervical cancer, had advanced disease at the time of diagnosis and had histopathologically confirmed squamous cell carcinomas. The low rates of literacy and knowledge regarding cervical cancer in this population were also associated with lower screening rates. Future interventions to address the cervical cancer burden in Ethiopia should include an effective educational component which has been shown to increase screening rates and ultimately decrease the cervical cancer incidence.

## Acknowledgements

We would like to acknowledge the Jimma University Teaching Hospital and the Global Physicians Corps for their financial and technical support in this study, and to the Touro University California Institutional Review Board in the United States of America, the Research and Publication Committee of the Faculty of Medical Sciences at Jimma University, the Jimma University Ethics Review Committee and the Jimma University Teaching Hospital Departments of Obstetrics and Gynecology and Medical Laboratory Sciences and Pathology in Ethiopia for their approval and permission to perform this study. We would also like to acknowledge all of the physicians/trainees/staff who assisted in data collection and to all of the study participants who provided this vital data in an overall effort to study and reduce the morbidity/mortality attributed to cervical cancer.

